# Myeloid-derived suppressor cells exacerbate poly(I:C)-induced lung inflammation in mice with renal injury and older mice

**DOI:** 10.1101/2023.06.07.544030

**Authors:** Zhiqi Xie, Haoyang Zhou, Masanori Obana, Yasushi Fujio, Naoki Okada, Masashi Tachibana

## Abstract

Viral pneumonia is a global health burden with a high mortality rate, especially in the elderly and in patients with underlying diseases. Recent studies have found that myeloid-derived suppressor cells (MDSCs) are abundant in these patient groups; however, their roles in the progression of viral pneumonia remain unclear. In this study, we observed a substantial increase in MDSCs in a mouse model of renal ischemia/reperfusion (I/R) injury and in older mice. When intranasal polyinosinic-polycytidylic acid (poly(I:C)) administration was used to mimic viral pneumonia, we found that mice with renal I/R injury exhibited more severe lung inflammation than sham mice when challenged with poly(I:C). In addition, MDSC depletion attenuated lung inflammation in mice with I/R injury. Similar results were obtained in older mice compared with those in young mice. Furthermore, we found that the adoptive transfer of *in vitro*-differentiated MDSCs exacerbated poly(I:C)-induced lung inflammation. Taken together, these experimental results suggest that the increased proportion of MDSCs in mice with renal I/R injury and in older mice exacerbates poly(I:C)-induced lung inflammation. These findings have important implications for the treatment and prevention of severe lung inflammation caused by viral pneumonia.

## Introduction

Viral lung infections are a considerable global health burden: in patients with highly pathogenic respiratory viral infections, pneumonia and the resulting acute respiratory distress syndrome, septic shock, and multiple organ failure are major risk factors for severe and fatal illnesses [1-3]. Multiple patient surveys from the 2003 severe acute respiratory syndrome (SARS) and coronavirus disease 2019 (COVID-19) outbreaks revealed that elderly patients, as well as those with underlying medical conditions such as kidney disease, diabetes, hypertension, cancer, and immunosuppression, are prone to developing severe illness [4-6]. Since these individuals are often immunocompromised, the immune response to antiviral therapy may lead to death due to complications related to the underlying disease. For example, patients with diabetes and hypertension have a sustained increase in pro-inflammatory cytokines caused by a dysregulated immune response, which may skew the antiviral response toward an inflammatory response associated with cytokine storms, tissue damage, and respiratory failure [7-9]. Older patients with chronic kidney disease, who are most at risk for death due to COVID-19, often have immune senescence and immunosuppressive conditions that hinder the approaches used to combat SARS-CoV-2 infection [10]. Unfortunately, the cascade of pathological and immune events and the key mechanisms involved in the aggravation of viral pneumonia remain unclear.

Myeloid-derived suppressor cells (MDSCs) are immune cells with suppressive functions that have received considerable attention in recent decades. MDSCs are a heterogeneous group of myeloid cells that can be classified as CD11b^+^Ly-6G and Ly-6C^hi^ monocytic MDSCs (M-MDSCs) and CD11b^+^Ly-6G^+^Ly-6C^int^ polymorphonuclear MDSCs (PMN-MDSCs) based on their morphology and surface marker expression in mice [11]. It has been reported that M-MDSCs suppress T cell proliferation via arginase 1 and inducible nitric oxide synthase (iNOS), whereas PMN-MDSCs exert inhibitory effects through arginase 1 and reactive oxygen species (ROS) [12]. Although MDSCs were originally described in patients with cancer, recent studies have highlighted their important roles in regulating immune responses in other pathological conditions, including infection, transplantation, and autoimmune diseases [13, 14]. In addition, there is compelling evidence that aging increases the number of circulating MDSCs in humans and mice [15], while CD11b^+^Gr-1^+^ cells isolated from the spleens of older mice can effectively inhibit T cell proliferation and activity [16]. Other studies have shown that MDSCs can exert suppressive effects through various pathways that ameliorate acute kidney injury or diabetic kidney disease [17, 18]. Together, these observations suggest that MDSCs may prevent excessive inflammation caused by aging and kidney disease. Paradoxically, MDSCs can also promote tissue degeneration and increase the risk of infection complications [19, 20], and exacerbate kidney damage [21]. Furthermore, MDSCs can display features of pro-inflammatory cells and contribute toward hyperinflammation under certain conditions [22-24], suggesting that the behavior of MDSCs depends on the context of the disease.

Several studies have highlighted the potential role of MDSCs in viral infections, including influenza A virus (IAV), hepatitis C virus (HCV), and SARS-CoV-2 [25-28]. In patients with COVID-19, it has been reported that MDSC expansion after infection correlates with disease severity and mortality [24]. Available data also suggests a direct role for MDSCs in exacerbating respiratory viral infections [27]. Since increased morbidity and mortality rates are consistently observed in aging individuals and in those with chronic diseases during viral infections, the increased frequency of MDSCs in such individuals may play a detrimental role in the progression of viral pneumonia. In this study, older mice and those with renal ischemia/reperfusion (I/R) injury were challenged with polyinosinic-polycytidylic acid (poly(I:C)) as a viral RNA analog to induce lung inflammation. Together, our experimental results reveal the role of MDSCs in these models of poly(I:C)-induced lung inflammation and have important implications for the treatment and prevention of severe lung inflammation caused by viral pneumonia in aging individuals and in those with chronic diseases.

## Materials and Methods

### Mice

Inbred male C57BL/6J mice (6-8 weeks old) were purchased from Japan SLC (Shizuoka, Japan) and were aged over 24 weeks in our facility. All animals were bred and maintained under pathogen-free conditions.

### Preparation of poly(I:C)

High-molecular-weight poly(I:C) (InvivoGen, CA, USA) was prepared according to the manufacturer’s instructions. Briefly, endotoxin-free water (provided by the manufacturer) was added to poly(I:C) to produce a final concentration of 4 mg/mL, incubated in a hot water bath (65-70 °C) for 10 min, and allowed to cool slowly to room temperature (about 25°C) to ensure proper annealing. The poly (I:C) solution was then aliquoted and stored at -20 °C. Before use, the poly(I:C) solution was diluted and vortexed to ensure thorough mixing.

### Murine model of poly(I:C)-induced pneumonia

Male C57BL/6J mice (6-8 weeks old; young mice, or over 24 weeks old; older mice) were anesthetized using isoflurane. Different doses (20, 50, and 100 μg) of poly(I:C) in 50 μL sterile PBS or PBS alone were administered intranasally (*i*.*n*.) twice through both nostrils alternately [41, 42]. Mice received seven poly(I:C) (or PBS) administrations, with a 24 h rest period between each administration. Anti-Ly-6G (clone1A8, 2 mg/kg; BioXCell, NH, USA) and anti-Ly-6C (clone: Monts 1, 2 mg/kg; BioXCell) were administered by intraperitoneal (*i*.*p*.) injection one day before poly(I:C) challenge, with one more dose injected after three days. Mice were sacrificed 7 days after poly(I:C) injection and retro-orbital blood (approximately 75 μL) was collected for flow cytometry analysis. Blood cell counts were determined using an XT-2000i automated hematology analyzer (Sysmex, Kobe, Japan). Lungs and BALF were obtained for further analysis.

### Murine model of renal I/R injury

The mouse model of renal I/R injury was established as described previously [43, 44]. Briefly, 6-8 weeks old male mice were anesthetized using isoflurane. A left unilateral flank incision was made and renal pedicle dissection was performed. A microvascular clamp (Natsume Seisakusho, Japan) was placed on the renal pedicle for 22 min while the animal was kept at a constant temperature and adequately hydrated. The clamp was then removed, the wound was sutured, and the mice were allowed to recover. After seven days, retro-orbital blood or spleen samples were collected to evaluate MDSC levels. Anti-Ly-6C and Anti-Ly-6G antibodies and poly(I:C) were administered.

### Bronchoalveolar lavage fluid (BALF) collection

After mice had been sacrificed, an abdominal cavity incision was made and a microvascular clamp was placed in the bronchus of the left lung. BALF was obtained by inserting a 20-gauge catheter into the trachea, through which 0.5 mL of cold Hank’s Balanced Salt Solution (HBSS; Gibco, USA) was flushed back and forth three times. BALF was centrifuged at 330 × *g* for 5 min at 4 °C. Cell-free supernatants were used to measure cytokine concentrations using Bio-Plex. The BALF cell pellet was treated with red cell lysis buffer and re-suspended in Hank’s Balanced Salt Solution (HBSS) supplemented with 2% fetal bovine serum (FBS) (2% FBS/HBSS; Gibco, CA, USA) for cell counting and flow cytometry analysis.

### Quantitative reverse transcription polymerase chain reaction (qRT-PCR)

Total RNA was isolated from CD11b^+^Gr-1^+^ cells purified from murine splenocytes using a JSAN cell sorting instrument (KS-Techno, Chiba, Japan) with TRIzol reagent. cDNA was synthesized using a QuantiTect reverse transcription kit (Qiagen, Hilden, Germany) according to the manufacturer’s instructions. qRT-PCR was performed using SYBR Premix Ex Taq (Tli RNaseH Plus; TaKaRa, Tokyo, Japan) on a CFX96 Touch Real-Time PCR Detection System (Bio-Rad, CA, USA). The specific primer sequences used are listed in Table S1. Glyceraldehyde 3-phosphate dehydrogenase (*Gapdh*) was used as a reference gene and the relative expression of other genes was calculated using the 2^-ΔΔCt^ method.

### Flow cytometry analysis

Cells were pelleted, washed with 2% FBS/HBSS, blocked with TruStain fcX (anti-mouse CD16/32) antibodies (BioLegend, CA, USA) for 5 min, and then stained with the following antibodies for 15 min at 4 °C: APC anti-mouse CD11b, Pacific Blue anti-mouse Gr-1, APC-Cy7 anti-mouse Ly-6C, FITC anti-mouse Ly-6G, APC anti-mouse CD3ɛ, Pacific Blue anti-mouse CD4, PE anti-mouse NK1.1, and FITC anti-mouse CD8α (BioLegend). The cells were then washed and resuspended in 2% FBS/HBSS. Shortly before performing measurements, a 7-amino actinomycin D viability staining solution (BioLegend) was added to each sample to stain dead cells. Flow cytometry analysis was performed using a BD FACSCanto II flow cytometer (BD Biosciences, NJ, USA). Data were analyzed using FlowJo software (version 10.7.0, BD Biosciences). The gating strategy used for flow cytometry analysis is as follows: monocytes (7AAD^-^CD45^+^CD11b^+^Ly-6G^-^Ly-6C^hi^), neutrophils (7AAD^-^CD45^+^CD11b^+^Ly-6G^+^Ly-6C^int^), CD4^+^ T cells (7AAD^-^CD45^+^CD3ɛ^+^CD4^+^NK1.1^-^), CD8^+^ T cells (7AAD^-^CD45^+^CD3ɛ^+^CD8α^+^NK1.1^-^) and NK cells (7AAD^-^ CD45^+^CD3ɛ^-^NK1.1^+^) (Fig. S1A).

### Histopathological examination

The left lungs were removed from euthanized mice, fixed in 10% formalin, and sent to the Kyoto Institute of Nutrition & Pathology for paraffin embedding. Whole lungs were cut into 4 μm sections, stained with hematoxylin and eosin (H&E), and imaged using the SLIDEVIEW VS200 Imaging System (EVIDENT, Tokyo, Japan). Digital images were imported into HALO software (Indica Labs) for analysis. Regions of interest around the relevant areas in each slide were annotated manually and lung sections were divided into normal and inflamed areas using the Indica Labs’ Area Quantification module (Version 1.0). Nuclear cells in the inflamed areas (infiltrating inflammatory cells) were automatically counted using the CytoNuclear v2.0.9 analysis module (Fig. S1B).

### Bio-Plex cytokine analysis

To detect multiple cytokines in BALF, the Bio-Plex Pro mouse cytokine assay (23-Plex Group I; Bio-Rad) was performed using a Luminex-xMAP/Bio-Plex 200 System with Bio-Plex Manager 6.2 software (Bio-Rad). Cytokine levels were measured using a cytometric magnetic bead-based assay according to the manufacturer’s instructions.

### Statistics

Significant differences were assessed using the Student’s *t* test or one-way analysis of variance (ANOVA) in GraphPad Prism (GraphPad Software). *P* values < 0.05 were considered statistically significant.

## Results

### MDSCs aggravate poly(I:C)-induced lung inflammation in mice with renal I/R injury

Poly(I:C), a synthetic analog of double-stranded RNA, is present in some viruses and is therefore widely used to model viral pneumonia [29]. Upon binding to toll-like receptor 3 (TLR3), retinoic acid-inducible gene I protein (RIG-I), melanoma differentiation-associated gene 5 (MDA5), and poly(I:C) selectively activate innate immune signaling pathways leading to inflammation [30]. First, we aimed to elucidate the importance of MDSCs in poly(I:C)-induced lung inflammation using a mouse model of acute renal I/R injury. A substantial increase in both MDSC subsets was observed in mice with renal I/R injury (Fig. 1A) and CD11b^+^Gr-1^+^ MDSCs showed increased expression of the immunosuppression-associated genes, *Arg1, Nos2*, and *Cybb* (Fig. 1B).

**Figure 1.**
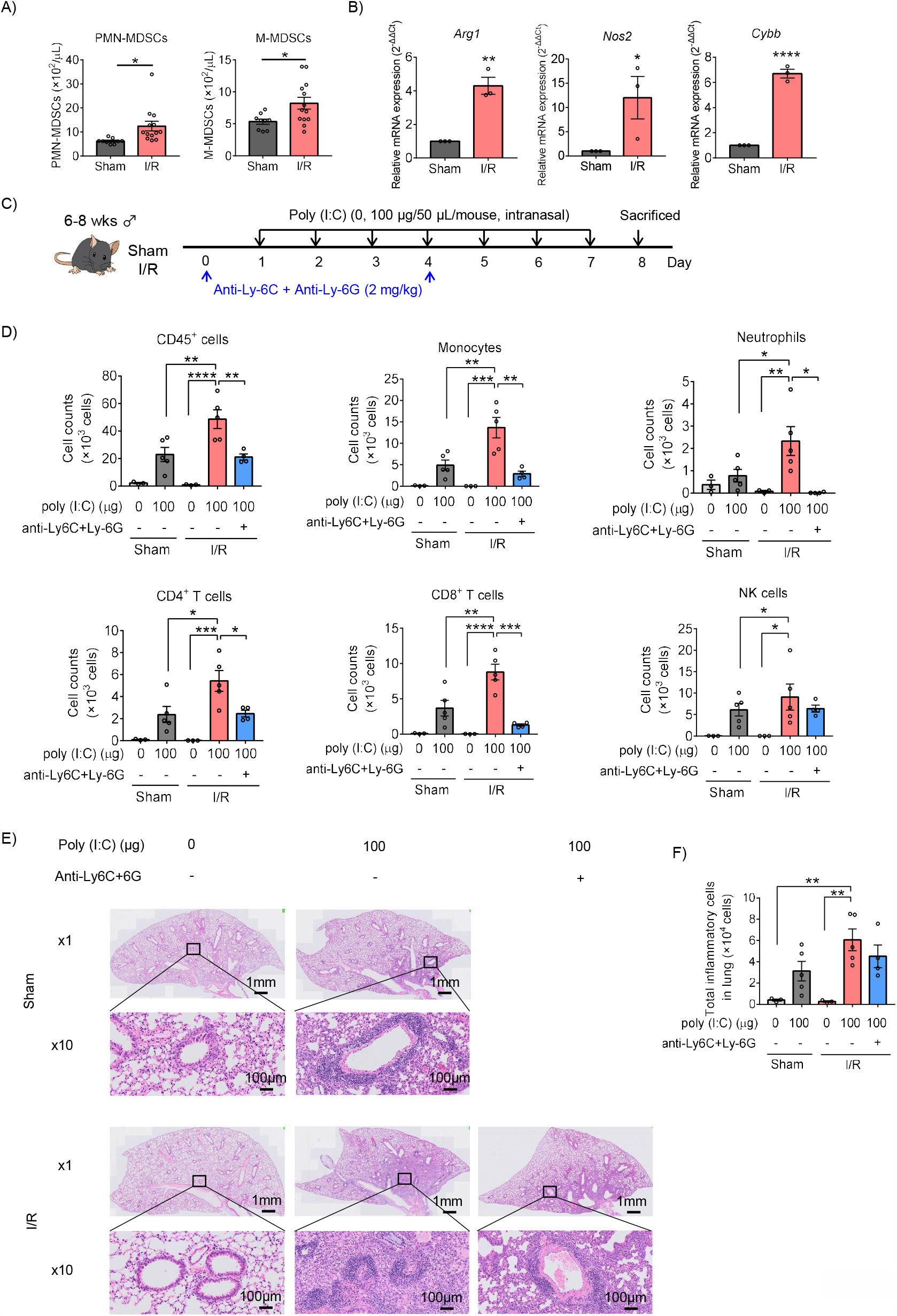
MDSCs aggravate poly(I:C)-induced lung inflammation in mice with renal I/R injury. A) MDSC subsets in the blood were counted (mean ± SEM of two independent experiments; *n* = 9 in sham group, *n* = 13 in I/R group. Student’s *t* test: \**p* < 0.05). B) *Arg1, Nos2*, and *Cybb* mRNA expression in MDSCs (CD11b^+^Gr-1^+^) sorted from sham or I/R injury mouse spleens measured using qRT-PCR (mean ± SEM; *n* = 3 per group, Student’s *t* test: \**p* < 0.05). C) Mouse model of poly(I:C)- induced pneumonia established using an anti-Ly-6C/Ly-6G dosing schedule. D) Total number of CD45^+^ cells, monocytes, neutrophils, CD4^+^ T cells, CD8^+^ T cells, and NK cells in BALF assessed using flow cytometry (mean ± SEM; one-way ANOVA: \**p* < 0.05, \*\**p* < 0.01, \*\*\**p* < 0.001, \*\*\*\**p* < 0.0001). E) Representative H&E-stained lung sections from Sham, I/R, and MDSC-depleted I/R groups at 1× and 10×. F) Total inflammatory cells in H&E-stained lung sections identified using HALO AI (mean ± SEM; one-way ANOVA: \*\**p* < 0.01)

Repetitive intranasal administration with poly(I:C) significantly increased the number of CD45^+^ cells in BALF (Fig. 1C, D). The increase in total cellularity in BALF was caused by significant monocyte, neutrophil, CD4^+^ T cell, CD8^+^ T cell, and NK cell infiltration (Fig. 1D). Notably, the number of inflammatory cells in BALF samples harvested from mice with I/R injury mice was significantly increased compared to sham mice.

To elucidate whether the effects induced by poly(I:C) in I/R-injured mice depended on MDSCs, I/R-injured mice were treated with anti-Ly-6C and anti-Ly-6G antibodies to deplete circulating MDSCs. Almost all M-MDSCs and total PMN-MDSCs were depleted from the blood (Fig. S2). In addition, MDSC-depleted I/R-injured mice displayed reduced inflammatory cell infiltration, especially for monocytes, neutrophils, CD4^+^ T cells, and CD8^+^ T cells (Fig. 1D).

Histological analysis of the lungs was performed to better understand the pathology induced by poly(I:C). Marked perivascular and moderate peribronchiolar interstitial inflammatory infiltrate was observed in poly(I:C)-treated sham mice (Fig. 1E). Meanwhile, more severe inflammatory infiltrate was observed in I/R injured mice and the inflammatory infiltrate in I/R-injured mice was slightly lower under MDSC-depleted conditions (Fig. 1E, F). These results suggest that the frequency of MDSCs is increased in mice with renal I/R injury and aggravates poly(I:C)-induced lung inflammation.

### MDSCs aggravate poly(I:C)-induced lung inflammation in older mice

Next, we assessed the role of MDSCs in poly(I:C)-induced lung inflammation in older mice. Consistent with previous reports, older mice showed an increase in both MDSC subsets compared to young mice (Fig. 2A). In addition, CD11b^+^Gr-1^+^ MDSCs isolated from older mice showed increased *Arg1, Nos2*, and *Cybb* expression compared to those isolated from young mice (Fig. 2B). Older mice also showed a significant increase in the number of CD45^+^ cells in BALF compared to young mice with poly(I:C) challenge, similar to mice with I/R injury (Fig. 2C, D). This increase in total cellularity in BALF samples from older mice was caused by significant neutrophil, CD4^+^ T cell, and CD8^+^ T cell infiltration. Conversely, MDSC depletion significantly decreased the levels of all analyzed cells in older mice (Fig. 2D). No significant reduction in total CD45^+^ cells was observed in young mice, only a decrease in neutrophils and NK cells (Fig. 2D). Analysis of lung peribronchial and perivascular inflammatory cells from lung sections revealed more severe inflammatory infiltrate in older mice than in young mice (Fig. 2E, F), while MDSC depletion reduced poly(I:C)-induced lung inflammation in older mice to the same level as in young mice. Together, these results suggest that MDSCs are upregulated in older mice and aggravate poly(I:C)-induced lung inflammation.

**Figure 2.**
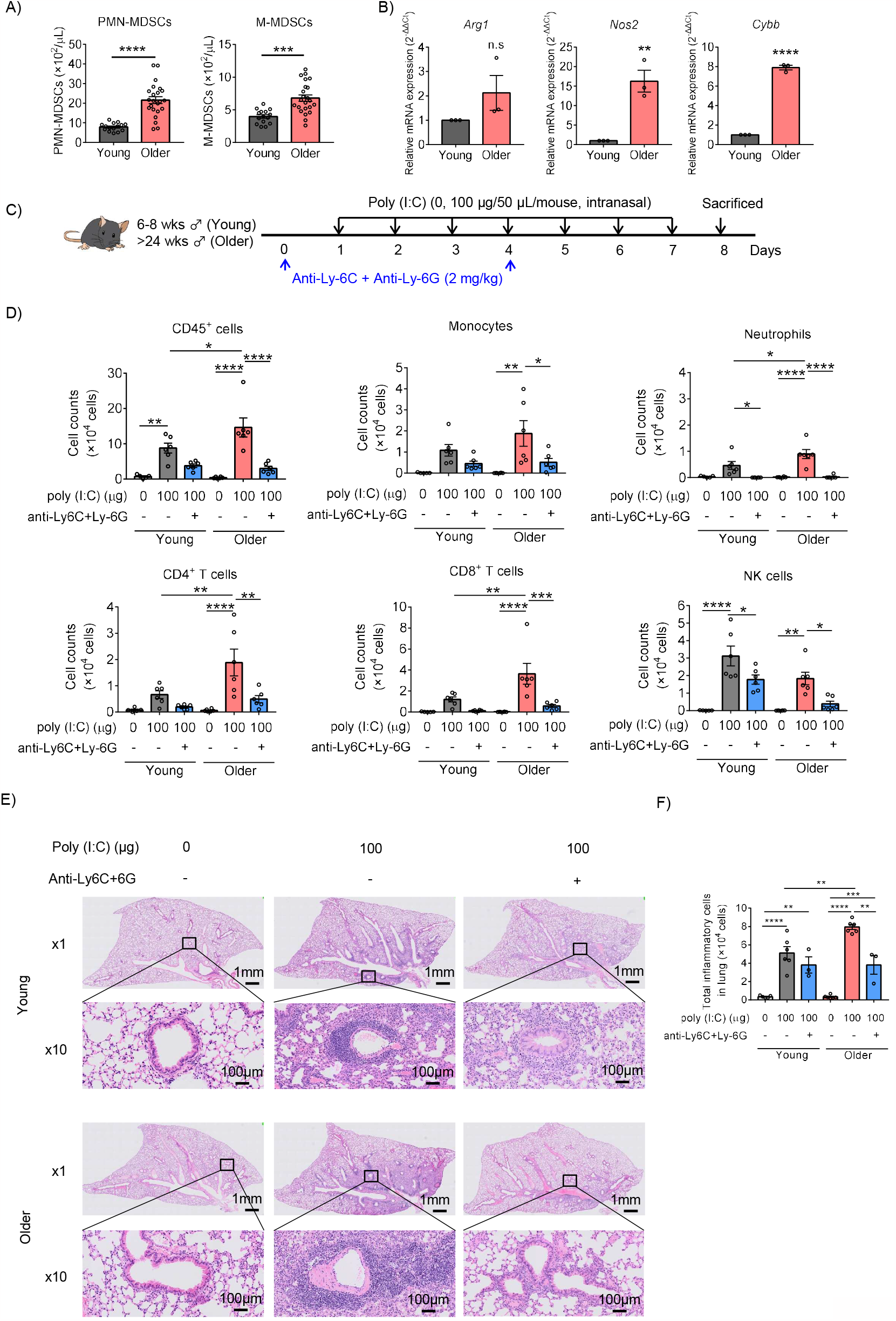
MDSCs aggravate poly(I:C)-induced lung inflammation in older mice. A) MDSC subsets in the blood were counted (mean ± SEM of two independent experiments; *n* = 15 in young, *n* = 23 in older. Student’s *t* test: \**p* < 0.05). B) *Arg1, Nos2*, and *Cybb* mRNA expression in MDSCs (CD11b^+^Gr-1^+^) sorted from young or older mouse spleens measured using qRT-PCR (mean ± SEM; *n* = 3 per group. Student’s *t* test: \**p* < 0.05). C) Mouse model of poly(I:C)-induced pneumonia was established and treated with anti-Ly-6C/Ly-6G antibodies per the dosing schedule shown. D) Total number of CD45^+^ cells, monocytes, neutrophils, CD4^+^ T cells, CD8^+^ T cells, and NK cells in BALF assessed using flow cytometry (mean ± SEM; one-way ANOVA: \**p* < 0.05, \*\**p* < 0.01, \*\*\**p* < 0.001, \*\*\*\**p* < 0.0001). E) Representative H&E-stained lung sections from sham, I/R, and MDSC-depleted I/R groups at 1× and 10×. F) Total inflammatory cells in H&E-stained lung sections identified using HALO AI (mean ± SEM; one-way ANOVA: \**p* < 0.05, \*\**p* < 0.01, \*\*\**p* < 0.001, \*\*\*\**p* < 0.0001)

### Adoptive transfer of MDSCs aggravates poly(I:C)-induced lung inflammation

To verify the direct effect of MDSCs on poly(I:C)-induced lung inflammation, *in vitro* MDSCs were adoptively transferred into mice in an intravenous manner and the consequences of inflammation were studied. *In vitro* MDSCs were differentiated from BM cells with GM-CSF stimulation, as described previously (CD11b^+^Gr-1^+^ MDSC purity over 90%, Fig. S3A) [31, 32]. These cells displayed higher *Arg1, Nos2*, and *Cybb* expression than BM cells (Fig. S3B) and potently inhibited CD8^+^ T cell proliferation (Fig. S3C).

Intranasal poly(I:C) administration dose-dependently increased inflammatory cell infiltration into the lungs in both the phosphate buffer saline (PBS) and MDSC transfer groups. The adoptive transfer of MDSCs significantly increased BALF CD45^+^ cell numbers, even in the absence of poly(I:C), mainly due to CD4^+^ T cell and neutrophil infiltration into the lung. When poly(I:C) was administered, monocyte and CD8^+^ T cell infiltration also increased. Analysis of lung peribronchial and perivascular inflammatory cells from lung sections revealed more severe lung inflammation after the adoptive transfer of *in vitro* MDSCs (Fig. 3C and D). Multiple poly(I:C) administrations upregulated the pro-inflammatory cytokines GM-CSF, IFN-α, IL-10, MCP-1, and TNF-α in BALF, which were further upregulated by the adoptive transfer of *in vitro* MDSCs (Fig. 3E). Thus, MDSCs appear to aggravate poly(I:C)-induced lung inflammation.

**Figure 3.**
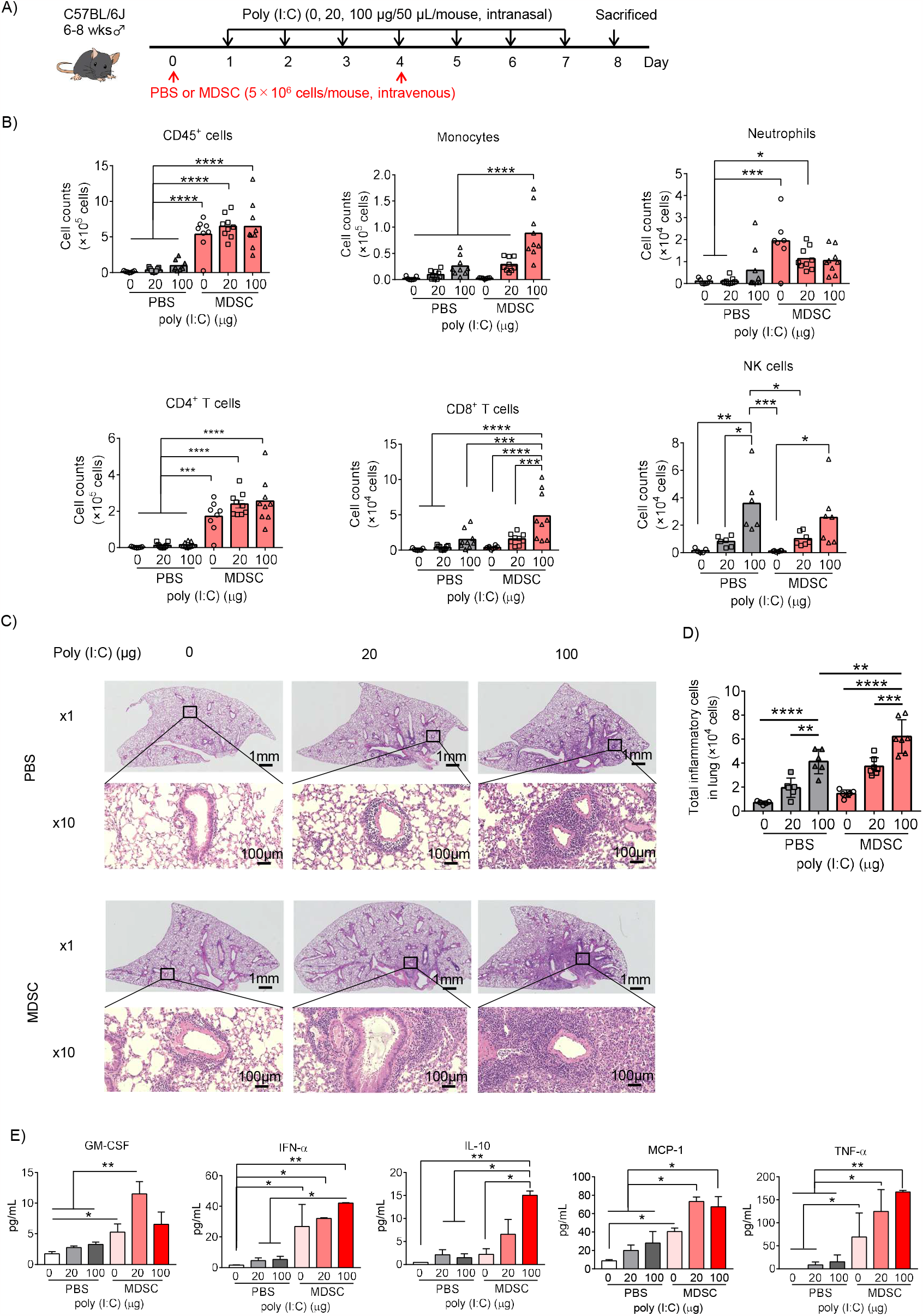
Adoptively transferred MDSCs aggravate poly(I:C)-induced lung inflammation. A) Mouse model of poly(I:C)-induced pneumonia established as shown with MDSC adoptive transfer. B) Total number of CD45^+^ cells, monocytes, neutrophils, CD4^+^ T cells, CD8^+^ T cells, and NK cells in BALF assessed using flow cytometry (mean ± SEM; one-way ANOVA: \**p* < 0.05, \*\**p* < 0.01, \*\*\**p* < 0.001, \*\*\*\**p* < 0.0001). C) Representative H&E-stained lung sections from PBS or MDSC-transferred mice at 1× and 10×. D) Total inflammatory cells in H&E-stained lung sections identified using HALO AI (mean ± SEM; one-way ANOVA: \*\**p* < 0.01, \*\*\**p* < 0.001). E) Cytokines in BALF were analyzed using Bio- Plex (mean ± SEM; one-way ANOVA: \**p* < 0.05, \*\**p* < 0.01).

## Discussion/Conclusion

Viral infections target the airway and alveolar epithelial cells, causing alveolar epithelial injury that can lead to acute respiratory distress syndrome and even death [2, 33]. Although dysregulated immune responses are hallmarks of severe infectious diseases, it remains unclear which innate and adaptive immune cells are critically involved in disease pathogenesis and which immunological mechanisms could be useful therapeutic targets. Higher morbidity and mortality rates in the elderly and in patients with chronic diseases may be related to increased MDSC levels. In this study, we showed that the frequency of MDSCs is increased before infection in older mice and in those with renal I/R injury, and is involved in the progression of viral pneumonia, leading to increased lung inflammation.

MDSCs exert immunosuppressive functions that may increase disease severity and cause clinical deterioration in patients with infectious diseases. However, MDSCs may also display features of pro-inflammatory cells that contribute toward hyper-inflammation under certain conditions [22-24]. Various factors secreted by inflamed tissues, such as GM-CSF, can promote the local recruitment of MDSCs from the circulatory system and promote their terminal differentiation into mature myeloid cells, as well as their activation to a pro-inflammatory phenotype with enhanced cytokine secretory capacity [34]. These studies may explain the significant increase in the inflammatory cytokines GM-CSF, IFN-α, MCP-1, and TNF-α observed in our study following the adoptive transfer of *in vitro* MDSCs in the absence of poly(I:C) challenge, suggesting that MDSCs may be a source of these cytokines. Consequently, MDSCs are probably dominant pathogenic factors in infectious diseases that drive exaggerated inflammation and the migration of immune cells into the lung.

In older mice, all inflammatory cells analyzed in this study showed high infiltration in BALF when poly(I:C) was administered, whereas their infiltration was decreased by the depletion of Ly-6C^+^ and Ly-6G^+^ cells. Thus, Ly-6C^+^ and/or Ly-6G^+^ cells may be the key causes of inflammatory cell infiltration into the lungs of older individuals. In I/R-injured mice, NK cell infiltration was increased following the depletion of Ly-6C^+^ and Ly-6G^+^ cells compared to other cells. Consistently, NK cell levels did not increase significantly with the adoptive transfer of *in vitro* MDSCs. I/R-injured mice were almost the same age as the young mice, suggesting that NK cell infiltration is independent of MDSCs in young mice with or without disease. A massive increase in CD4^+^ T cells and neutrophils was observed after the adoptive transfer of *in vitro* MDSCs; however, this change did not exacerbate pneumonia in the absence of poly(I:C), indicating that CD8^+^ T cells, monocytes, and NK cells may play roles in the exacerbation of pneumonia. Recent studies have demonstrated that NK cells exert anti-SARS-CoV-2 activity but show defects in viral control, cytokine production, and cell-mediated cytotoxicity in patients with severe COVID-19 [35-37]. When combined with our results, these findings suggest that NK cells are a key factor in the deterioration of patients with pneumonia, in addition to MDSCs.

Since we found that MDSC depletion reduces poly(I:C)-induced lung inflammation in older mice and those with I/R injury, our study highlights the potential of therapeutic approaches that aim to reduce the number of MDSCs. Preliminary studies have shown that a CCR5 inhibitor can alleviate SARS-CoV-2 plasma viremia in patients with COVID-19 [38]. Targeting the CCL5/CCR5 axis can reduce the recruitment of MDSCs from the bone marrow to the lesion site [39]; thus, COVID-19-related immunomodulatory disorders could be improved by targeting this pathway. The absence of HLA-DR is an important marker of human MDSCs and one study has shown that IL-6 blockers can partially elevate HLA-DR expression, considering the decreased MDSC levels in patients with severe COVID-19 [40]. Taken together, these data suggest that new approaches targeting MDSCs could be used to treat and prevent severe lung inflammation caused by COVID-19. It should be noted that although MDSC depletion in older mice reduced the severity of pneumonia to levels observed in young mice, MDSC depletion in young mice did not significantly improve pneumonia. Therefore, MDSC depletion may only attenuate the worsening of pneumonia in elderly patients or in those with underlying diseases.

Research on the relationships between MDSCs and acute or chronic viral infections is still in its infancy and has only begun to gain widespread attention since the COVID-19 pandemic. For instance, several recent studies have demonstrated an increase in the frequency of MDSCs in patients with COVID-19, which is related to immune regulation during infection and can be used as an indicator of the severity of COVID-19 [24]. To our knowledge, no studies have yet reported the role of MDSCs in the progression of viral pneumonia in aging individuals and in those with chronic diseases. Although the details of the immune events and key mechanisms remain unclear, this study presents evidence that the increased MDSC profile present in older mice and those with renal I/R injury exacerbates poly(I:C)-induced lung inflammation. In addition, we demonstrated that adoptively transferred MDSCs could worsen poly(I:C)-induced lung inflammation, indicating that MDSCs play a direct role in the pathogenesis of pneumonia. It should be noted that poly(I:C) does not effectively mimic the viral replication process; therefore, the effect of MDSCs on viral clearance requires further investigation. Future studies should continue to investigate the role of MDSCs in viral progression and whether they could be a potential target for drug intervention in virus-infected mice or patients.

## Statement of Ethics

This study protocol was reviewed and approved by [The Animal Experiment Committee of Osaka University], approval number [Douyaku R03-7-2].

## Author Contributions

Z.X. designed experiments and wrote the manuscript. Z.X. performed experiments and H.Z. assisted in data acquisition. M.O. and Y. F. contributed toward the renal I/R injury model. N.O. helped with experimental design and assisted with discussions. M.T. conceptualized and supervised the study and wrote the manuscript. All authors helped to review and revise the manuscript.

## Data Availability Statement

The corresponding author will provide all the data used in this study upon request.

