## Supplemental Information for "Myeloid-derived suppressor cells exacerbate poly(I:C)-induced lung inflammation in mice with renal injury and older mice"

**This PDF file includes:**

Supplemental Methods
Figures S1 to S3
Table S1

**Supplemental Methods:**

***MDSC differentiation in vitro***

The *in vitro* differentiation of bone marrow (BM) cells into MDSCs (*in vitro* MDSCs) was performed as described previously (1, 2). Briefly, BM cells from C57BL/6J mice were stimulated with 40 ng/mL recombinant granulocyte-macrophage CSF (GM-CSF) (Peprotech, NJ, USA) for 4 days.

***In vitro suppression assay***

$CD8^+$ T cells were isolated from the spleens of C57BL/6J mice using the MojoSort magnetic cell separation system, as described previously (1, 2), and labeled with eFluor 670 proliferation dye (eBioscience, Thermo Fisher Scientific, CA, USA). eFluor 670-labeled $CD8^+$ T cells were incubated with *in vitro* differentiated MDSCs at different ratios in a 96-well plate cultured with anti-mouse CD3ε antibody/anti-mouse CD28 antibody (BioLegend). After three days of incubation at 37 °C in 5% $CO_2$, the proliferation of $CD8^+$ T cells, as determined using eFluor 670 fluorescence intensity, was analyzed using flow cytometry.

**Table S1.** List of primer sequences used in qRT-PCR analysis

| Gene | Forward (5' to 3') | Reverse (5' to 3') |
|---|---|---|
| *Gapdh* | TGACCTCAACTACATGGTCTACA | CCGTGAGTGGAGTCATACTGG |
| *Arg1* | CCTATGTGTCATTTGGGTGGATG | GGTTGTCAGGGGAGTGTTGAT |
| *Nos2* | GGAGTGACGGCAAACATGACT | TAGCCAGCGTACCGGATGA |
| *Cybb* | CCTCTACCAAAACCATTCGGAG | CTGTCCACGTACCGGATGA |

qRT-PCR, quantitative reverse transcription-polymerase chain reaction; Genes: *Gapdh*, glyceraldehyde 3-phosphate dehydrogenase; *Arg1*, arginase 1; *Nos2*, nitic oxide synthase 2; *Cybb*, cytochrome b-245 beta polypeptide

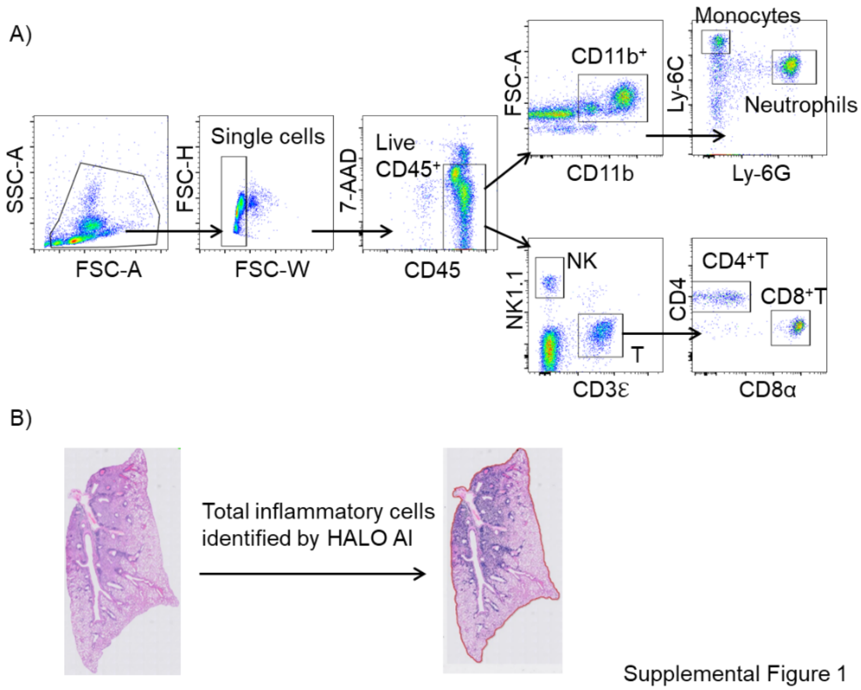


**Fig. S1.** A) Gating strategy used for flow cytometric analysis. Monocytes ($7AAD^-CD45^+CD11b^+Ly\text{-}6G^-Ly\text{-}6C^{hi}$), neutrophils ($7AAD^-CD45^+CD11b^+Ly\text{-}6G^+Ly\text{-}6\ C^{int}$), $CD4^+$ T cells ($7AAD^-CD45^+CD3\varepsilon^+CD4^+NK1.1^-$), $CD8^+$ T cells ($7AAD^-CD45^+CD3\varepsilon^+CD8\alpha^+NK1.1^-$), and NK cells ($7AAD^-CD45^+CD3\varepsilon^-NK1.1^+$). B) Representative image showing total inflammatory cells in H&E-stained sections identified using HALO AI

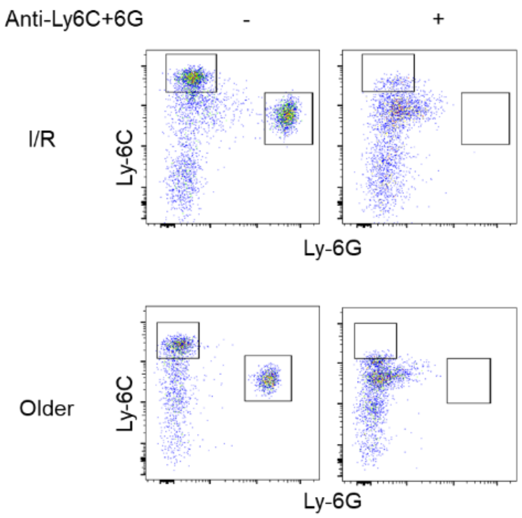


Supplemental Figure 2

**Fig. S2.** Representative flow cytometry analyses of the frequencies of M-MDSCs (CD11b$^{+}$Ly-6C$^{hi}$Ly-6G$^{-}$) and PMN-MDSCs (CD11b$^{+}$Ly-6 C$^{int}$Ly-6G$^{+}$) in blood from PBS- or anti-Ly-6C+anti-Ly-6G antibody-treated mice with I/R injury or older mice with poly(I:C) challenge prior to sacrifice

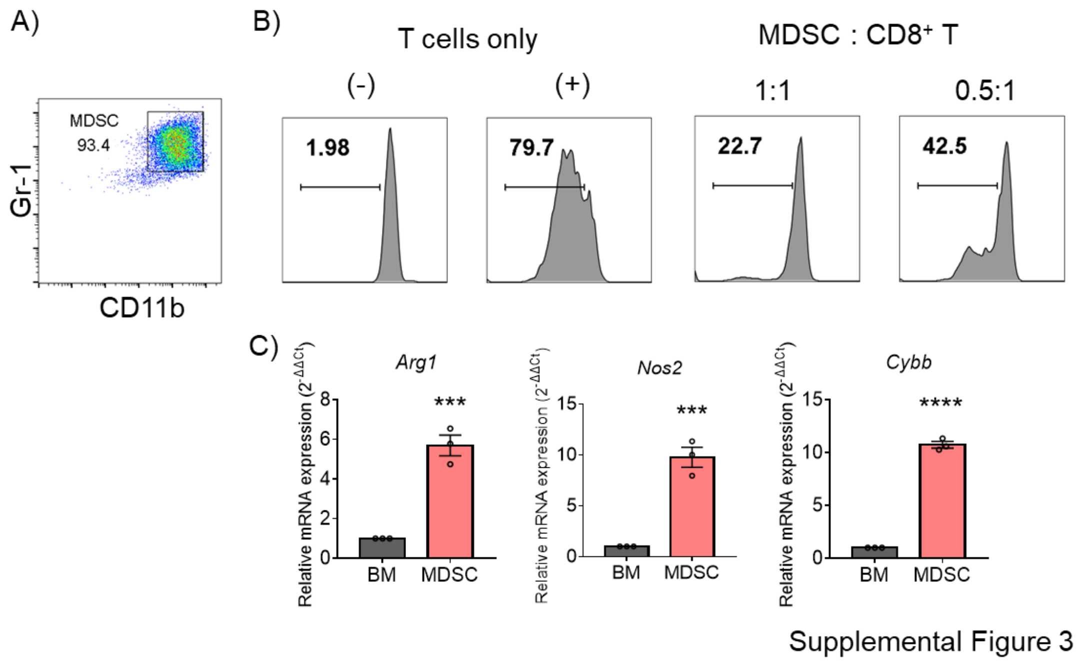


**Fig. S3.** A) Flow cytometry analysis of the percentage of CD11b$^+$Gr-1$^+$ populations after four days of culture in medium supplemented with GM-CSF (40 ng/mL). B) Histograms of eFluor 670 expression in CD8$^+$ T cells. MDSCs were combined at a 1:1 or 0.5:1 ratio with eFluor 670 labeled CD8$^+$ T cells, followed by stimulation with anti-CD3ε and anti-CD28 antibodies. (-) Mean CD8$^+$ T cells without stimulation with anti-CD3ε Ab and anti-CD28 antibodies. C) mRNA expression of *Arg1*, *Nos2*, and *Cybb* in BM cells or MDSCs (CD11b$^+$Gr-1$^+$) measured using qRT-PCR (mean ± SEM; $n$ = 3 per group. Student's $t$-test: *$p < 0.05$)
